# Loss-of-function coding variants in the Ras of Complex Proteins/GTPase domain of Leucine Rich Repeat Kinase 2

**DOI:** 10.1101/2024.12.07.627348

**Authors:** Sarah Butterfield, Susanne Herbst, Patrick Alfryn Lewis

## Abstract

The *LRRK2* gene is a key contributor to genetic risk of Parkinson’s disease, and a priority drug target for the disorder. Leucine Rich Repeat Kinase 2, the protein product of *LRRK2*, is a multidomain enzyme implicated in a range of cellular processes – including endolysosomal trafficking and damage response. Based on the report that truncation and structural variants resulting in loss of LRRK2 protein are observed in human populations, genomic sequence repositories were queried for coding variants affecting key catalytic residues in LRRK2 – resulting in the identification of three variants (K1374E, K1374R, and T1348P) predicted to ablate the capacity of LRRK2 to bind GTP. Biochemical and cellular characterization of these variants confirmed loss of GTP binding, as well as reduced or loss of kinase activity. These data demonstrate the presence of rare coding enzymatic loss-of-function variants in humans, with implications for our understanding of LRRK2 as a driver of disease and as a drug target.

## Introduction

Parkinson’s disease (PD) is a common neurodegenerative disease characterised by autonomic dysfunction, progressive movement disorder, and cognitive decline. This is coupled with neuronal loss and the accumulation of alpha synuclein in intracellular inclusions called Lewy bodies (1). Whilst idiopathic cases of multifactorial aetiology represent the majority of cases, around 5-10% of PD is associated with a familial pattern of inheritance (2). Of these familial cases, autosomal dominant coding mutations in the *LRRK2* gene on chromosome 12 are the most common cause, with more common coding and non-coding variants at the *LRRK2* locus also linked to increased lifetime risk of developing idiopathic Parkinson’s (3).

Leucine Rich Repeat Kinase 2 (LRRK2), the product of the *LRRK2* locus, is a multi-domain scaffold enzyme, which possesses both guanosine triphosphatase (GTPase) and kinase activities (4). The catalytic core of LRRK2 is composed of the ROCO/GTPase supradomain (a Ras of complex proteins [ROC] domain and a C-terminal of ROC [COR] domain), followed by a serine-threonine kinase domain. Notably, PD associated coding mutations in LRRK2 cluster within the enzymatic domains of the protein, and include the N1437H, R1441C (both in the ROC/GTPase domain), Y1699C (in the COR domain), and G2019S and I2020T mutations (both in the kinase domain). These variants act by distorting the enzymatic activities of LRRK2, increasing kinase activity and decreasing GTPase activity (5). LRRK2 has been reported to directly phosphorylate a number of substrates, in particular a subset of Rab GTPases. Mutations resulting in increased phosphorylation and disruption of intracellular trafficking – although the mechanisms connecting these events to neurodegeneration remain obscure (6). The prevalence of *LRRK2* mutations and the enzymatic activities of the protein have established LRRK2 as a leading target for drug discovery, with ongoing clinical trials for kinase inhibitors and antisense oligonucleotides (7).

One of the many intriguing aspects of the molecular genetics of *LRRK2* is the identification of loss-of-function (LOF) variants at the *LRRK2* locus (8). Initially identified through a PD case/control study, large scale analysis of the *LRRK2* locus from the Gnomad, 23andme, and UK Biobank cohorts revealed 1455 individuals with LRRK2 heterozygous LOF variants, including frameshifts, premature stop codons and alterations in splicing, with a population frequency of 0.48% (9). No evidence of association of heterozygous *LRRK2* LOF with disease has yet emerged, despite a significant reduction in LRRK2 protein levels.

Given the prominence of the enzymatic activities of LRRK2 in its function and disease association, the increasing comprehension of the structure/function of this protein and proliferation of exome sequence data across human populations provides an opportunity to test whether there are specific coding variants that result in loss of enzymatic function of LRRK2 - the subject of this current study.

## Materials and methods

### Cell culture

HEK293T cells were obtained from ATCC (ATCC CRL-3216; RRID:CVCL_0063) and cultured in DMEM/10% FCS. If HEK293T cells were seeded on for high-content imaging, the wells were pre-coated with Geltrex (# A1569601, ThermoFisher Scientific). All cells were cultured at 37 °C, 5% CO2.

### Plasmids and site-directed mutagenesis

The EGFP-LRRK2-WT plasmid (pcDNA5-FRT-TO-EGFP-LRRK2, DU13363) was purchased from the MRC PPU Reagents and Services, University of Dundee. LRRK2 mutant constructs were generated by site-directed mutagenesis using the Q5 site-directed mutagenesis kit from New England Biolabs (#E0552S, NEB). All plasmids were maintained in *E. coli* DH5α (#11583117, ThermoFisher Scientific), cultured at 30 °C, and extracted using the QIAprep Spin Miniprep Kit from Qiagen. Successful modification was verified by sequencing and newly extracted plasmids were verified by EcoRI digest.

### Transfection and stimulation

For WB, HEK293T cells were seeded in 12-well plates and incubated overnight. The cells were tranfected with 500 ng DNA using FuGENE HD Transfection Reagent (#E2311, Promega). For high-content imaging, cells were seeded in 96-well plates and reverse-transfected with 100 ng of plasmid DNA. For the GTP-pulldown, HEK293T cells were seeded in 60 mm dishes and transfected with 3 µg DNA. A DNA:FuGENE ratio of 1:3 was used for all transfections. To stimulate LRRK2 kinase activity, cells were treated with 1 mM L-leucyl-L-leucine methyl ester (LLOMe; BAChem, #4000725.0001) for 1 hr 24 hrs after transfection.

### GTP-agarose pull-down

Cells were lysed in cell lysis buffer (20 mM Tris–HCl, pH 7.4, 1% Triton X-100, 150 mM NaCl, 10% (v/v) glycerol, 1 mM EGTA) containing protease and phosphatase inhibitors by rotating at 4 °C for 45 min. The lysate was clarified by centrifuging at 4 °C, 16.6 x *g* for 10 min. 30 µl of the clarified lysate were removed and stored at –20 °C (“Input”). 30 µl of GTP-agarose beads (Sigma) were added to the remaining clarified lysate. The beads and lysate were incubated at 4 °C for 4 hrs on a rotating wheel. The beads were washed thrice with cell lysis buffer and bound proteins were eluted by incubating the beads with cell lysis buffer containing 10 mM GTP.

### Western Blotting

Samples were denatured at 80 °C for 8 minutes in LDS sample buffer containing 1x NuPAGE reducing agent (ThermoFisher Scientific). Samples were run on a NuPAGE 4-12 % Bis-Tris gel (ThermoFisher Scientific) and transferred to a PVDF membrane using a Turbo-Blot transfer system (Biorad). The PVDF membrane was blocked for 1 hour in a TBS-T-milk solution (5 % skimmed milk powder, TBS, 0.05 % Tween-20). After blocking, membranes were incubated at 4 °C overnight with primary antibodies diluted 1:1000 in TBS-T-milk, and then secondary antibodies diluted 1:10’000 in TBS-T-milk for 45 minutes at room temperature. Western blots were developed using an iBright imaging system (ThermoFisher Scientific) and quantified with densitometry using FIJI software (Schindelin et al. 2012). Primary antibodies used for Western blotting were rabbit-anti-LRRK2 (Abcam Cat# ab133474, RRID:AB_2713963), rabbit-anti LRRK2 pS935 (Abcam Cat# ab133450, RRID:AB_2732035), rabbit-anti-LRRK2 pS1292 (), rabbit-anti-Rab10 pT73 (Abcam Cat# ab241060, RRID:AB_2884876), rabbit-anti-Rab10 (Abcam Cat# ab237703, RRID:AB_2884879), rabbit-anti-Rab12 pS106 (Abcam Cat# ab256487, RRID:AB_2884880), rabbit-anti-Rab12 (Proteintech Cat# 18843-1-AP, RRID:AB_10603469) and mouse-anti-β-actin (Sigma-Aldrich Cat# A1978, RRID:AB_476692). Secondary antibodies were anti-rabbit-HRP (Sigma-Aldrich Cat# A0545, RRID:AB_257896) and anti-mouse-HRP (Sigma-Aldrich Cat# A3682, RRID:AB_258100).

### High-content imaging

Samples were fixed in 4 % methanol-free PFA (15710, Electron Microscopy Sciences) diluted in PBS for 15 min at 4°C. Permeabilisation and blocking was achieved by incubating samples in 0.3 % Triton-X100/5 % FCS/PBS for 20 min at room temperature. Primary antibodies were diluted 1:100 in the permeabilisation/blocking solution and incubated for 1 hr at room temperature. Secondary antibodies were diluted 1:1000 in the permeabilisation/blocking solution and incubated for 1 hr at room temperature. Nuclei were stained with DAPI (300 nM) for 10 min at room temperature and cells were imaged in PBS using the 63x objective on an OPERA Phenix high-content imager (PerkinElmer). Primary antibodies used for imaging were rabbit-anti-Rab10 pT73 (Abcam Cat# ab241060, RRID:AB_2884876) and mouse-anti-human-LAMP1 (DSHB Cat# h4a3, RRID:AB_2296838). Secondary antibodies were goat-anti-rabbit-AF568 (Thermo Fisher Scientific Cat# A-11011, RRID:AB_143157) and goat-anti-mouse-AF647 (Thermo Fisher Scientific Cat# A-21235, RRID:AB_2535804). Images were analysed for the % of Rab10 pT73 positive lysosomes using the Harmony software (PerkinElmer).

### Structural modelling

Residues involved in GDP-binding were identified in a cryo-EM structure of monomeric LRRK2 (PDB ID: 7LHW,(10)). To identify residues potentially involved in GTP-binding, LRRK2 (NP_940980.4) was modelled as a monomer containing GTP, Mg2+ and ATP as ligands using the AlphaFold3 server (11). All structures were displayed using Mol*Viewer (12).

## Results and discussion

### Naturally occurring LRRK2 GTP-binding site mutants lose the ability to bind GTP

The critical importance of residues K1347 and T1348 in the active site of the ROC domain of LRRK2 for GTP-binding and hydrolysis has previously been reported, with artificial mutations at these codons (K1347A and T1348N) ablating the capacity of LRRK2 to bind GTP (13, 14). The cryoEM derived GDP-bound structure of LRRK2 reveals that K1347 forms a hydrogen bond with the β-phosphate of GDP and T1348 forms a hydrogen bond with the α-phosphate (**Figure 1A**) (10). In accordance with the loss of GTP-binding of the K1347A and T1348N mutants, modelling GTP-binding using AlphaFold 3 predicts that residue K1347 directly interacts with the gamma-phosphate of GTP and that T1348 coordinates a Mg^2+^ that contributes to the positioning of the γ-phosphate (**Figure 1B**). To identify naturally occurring putative LRRK2 GTP-binding null mutants, the gnomAd and MCPS Variant Browser were accessed to search for variation at these residues, resulting in the identification of heterozygous carriers of K1347E, K1347R and T1348P variants (**Table S1**) (15, 16).

**Figure 1:**
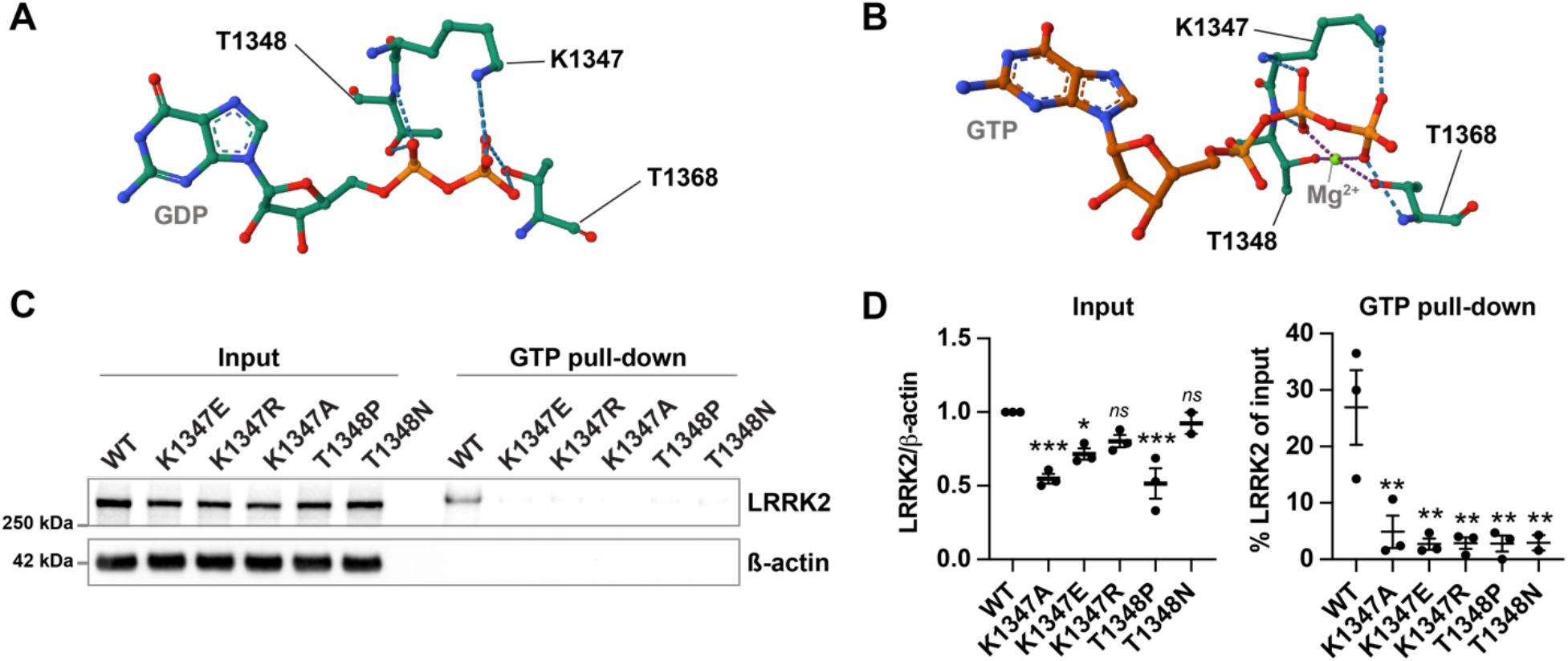
Naturally occurring LRRK2 GTP-binding variants lose the ability to bind GTP. (A)Contribution of K1347 (Lys1347) and T1348 (Thr1348) to GDP binding in the LRRK2 monomeric structure published by Myasnikov *et al*. (PDB 7LHW). (B)AlphaFold3 model of GTP-bound LRRK2 predicts coordination of the gamma-phosphate of GTP by K1347 and T1348. (C)LRRK2 GTP-agarose pull-down of artificial and naturally occurring K1347 and T1348 LRRK2 variants. HEK293 cells were transfected with the indicated LRRK2 constructs, and LRRK2 was pulled down from the cell lysates using GTP-agarose. WBs of the cell lysate (1 % of pull-down) and the pull-down were analysed by anti-LRRK2 staining. (D)Quantification of total LRRK2 levels from the input and the % of pulled down LRRK2. n=2-3. One-way ANOVA, followed by Dunnett’s multiple comparisons test against WT. *ns* – not significant, * *p > 0*.*05*, ** *p > 0*.*01*, *** *p > 0*.*001*.

To test the impact of these naturally occurring ROC domain variants on GTP-binding, these mutations were inserted into the full length LRRK2 open reading frame, alongside the previously characterised GTP-binding deficient artificial mutants K1347A and T1348N. These mutants displayed robust expression in HEK293 cells, although some of the GTP-binding variants displayed decreased expression levels when compared to WT (**Figure 1C, D**). GTP-binding was assessed by GTP-agarose bead pull-down. When compared to input levels, only WT LRRK2 was pulled down using GTP-agarose beads (**Figure 1C,D**), indicating that none of the naturally occurring LRRK2 GTP-binding site mutants bind GTP.

### Naturally occurring LRRK2 GTP-binding site mutations reduce kinase function

As the LRRK2 GTPase and kinase functions are highly interlinked, the impact of the LRRK2 GTP-binding site mutants on LRRK2 kinase activity was then investigated. The GTPase mutants were expressed in HEK293 cells, and LRRK2 kinase activity was assessed in the presence and absence of the lysotoxic agent LLOMe (which stimulates LRRK2 activity) by measuring the phosphorylation of the LRRK2 substrates Rab10 and Rab12, and LRRK2 autophosphorylation site S1292.

At both steady-state and following LLOMe stimulation, the K1347E and T1348P mutants displayed a marked reduction of kinase activity as evidenced by a loss in Rab10 and Rab12 phosphorylation, and a concomitant decrease in pS935 and pS1292 phosphorylation of LRRK2. In contrast, and surprisingly, the K1347R mutant retained partial kinase activity (**Figure 2A and B**). To test whether GTP-binding site mutant LRRK2 is recruited to lysosomes and is capable of activating Rab proteins following damage, cells expressing these variants were treated with LLOMe, and the response of LRRK2 assessed using fluorescence microscopy. Analysis of Rab10 pT73 activation and localization to lysosomes mirrored the results obtained by Western blot, with only the K1347R variant retaining Rab10 T73 phosphorylation (**Figure 2C and D**).

**Figure 2:**
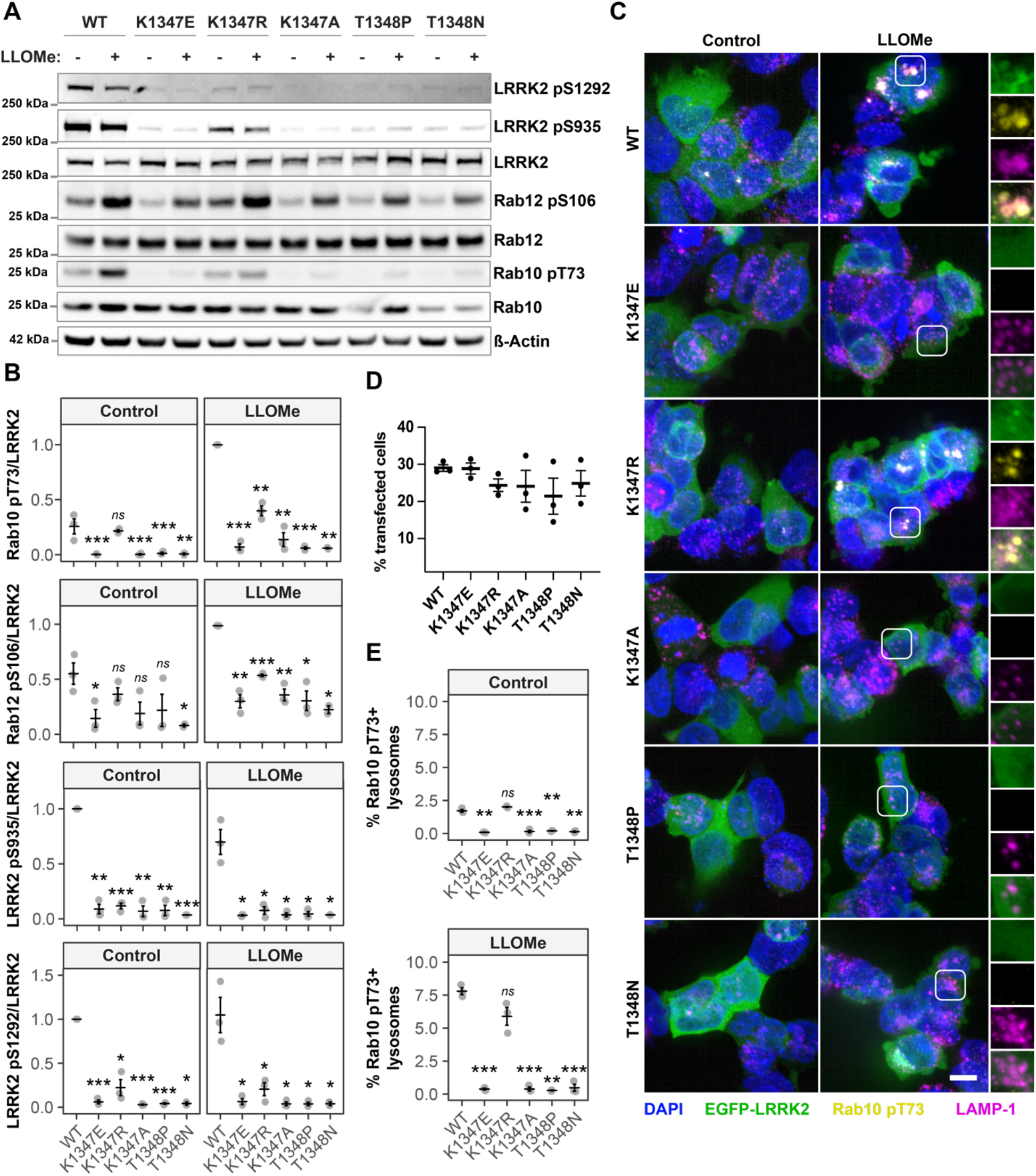
LRRK2 GTP-binding variants impair LRRK2 kinase activity. HEK293 cells were transfected with the indicated EGFP-LRRK2 GTP-binding variants and treated with 1 mM LLOMe for 1 hr where indicated. (A) LRRK2, Rab10 and Rab12 phosphorylation was assessed by Western blot. (B) Quantification of Rab10 T73 and Rab12 S106 phosphorylation (normalised to WT LLOMe) and LRRK2 S935 and S1292 phosphorylation (normalised to WT control) from (A). N=3. (C) Co-localisation of Rab10 pT73 with LAMP-1 positive lysosomes was assessed by high content imaging. Scale bar = 10 µm. (D) Quantification of the % of transfected cells from (C) n=3. (D) Quantification of the % of Rab10 pT73 positive lysosomes from (C) n=3. Statistical analysis was conducted by two-way anova followed by a t-test corrected for multiple comparisons against WT. *ns* – not significant, * *p > 0*.*05*, ** *p > 0*.*01*, *** *p > 0*.*001*.

Although it was not possible to directly quantify LRRK2 recruitment to lysosomes in this overexpression system, it was notable that the K1347 mutants were still recruited to lysosomes, whereas this was not the case for the T1348 mutants (**Figure S1**).

Taken together, these data suggest that naturally occurring variation at residues required for LRRK2 GTPase function causes a *de facto* loss of LRRK2 GTP hydrolysis function, resulting in ablation or a reduction in kinase activity. This expands the range and nature of loss-of-function variants observed in LRRK2, and highlights the need to consider coding variants as well as frameshift or premature stop codon mutations in the context of reduced LRRK2 activity.

No clinical phenotype has been reported with the K1347E, K1347R, or T1348P variants from the genomic repositories, and to date none of these have been identified in screens for disease associated mutations. Notably, the loss of enzymatic activity – while retaining scaffolding function – more closely matches the cellular consequences of treatment with small molecule LRRK2 kinase inhibitors compared to allelic variants resulting in complete loss of protein. Given that the majority of PD associated mutations result in increased LRRK2 kinase activity, it is unlikely that the ROC domain loss-of-function variants would predispose carriers to Parkinson’s, and may even reduce the risk of disease.

An intriguing, and perhaps perplexing, aspect of these results is the observation that the K1347R variant retained the ability to phosphorylate Rab substrates (albeit at a reduced level compared to WT), despite not being able to bind GTP-agarose. This discrepancy could be due to the GTP-agarose bead pull-down lacking sensitivity, or being limited to cellular lysates rather than performed in a living cellular environment, and not detecting residual GTP binding capacity for this variant. It is of note that the arginine side chain retains a positive charge, similar to the wild type lysine, although the increased size of the arginine side chain could result in steric hinderance of GTP binding.

In summary, this study reports the existence, and characterizes the consequences of, loss-of-function coding variants in the ROC/GTPase domain of LRRK2. This highlights the need to consider naturally occurring functional coding variants in the context of LRRK2 activity, and supports further investigation into whether these variants act to increase or decrease risk of disease.

## Acknowledgements

PAL is a Royal Society Industry research fellow in partnership with LifeArc technologies (IF\R2\222002). This research was funded in whole or in part [Grant numbers 18285 and 021184] by the Michael J. Fox Foundation for Parkinson’s Research (MJFF).

## Author CRediT statement

SB carried out the investigation and wrote the original draft

SH conceived, supervised and validated the experiments, and reviewed and edited the manuscript

PAL conceived and supervised the experiments, and reviewed and edited the manuscript.

