## Supplemental figure and table for "Loss-of-function coding variants in the Ras of Complex Proteins/GTPase domain of Leucine Rich Repeat Kinase 2"

Figure S1

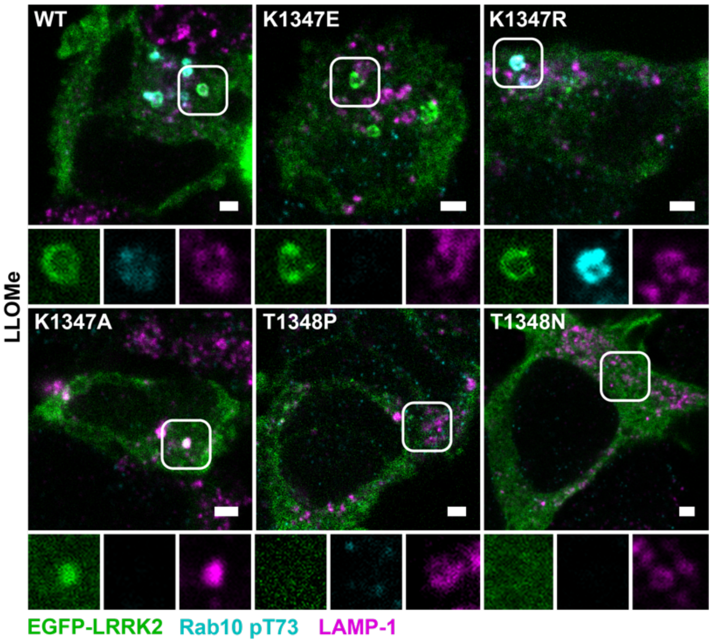


**Supplementary Figure 1: LRRK2 recruitment to lysosomes is impaired in the T1348 mutants.**

HEK293 cells were transfected with the indicated EGFP-LRRK2 constructs and treated with LLOMe (1 mM) for 1 hr. LRRK2 recruitment to LAMP1 positive lysosomes was assessed by confocal microscopy. Lysosomal recruitment was observed for the K1347E and K1347R mutants, but not detected for the T1348P and T1348N mutants.

| **Variant** | **Consequence** | **Allele Frequency** | **Source** |
|---|---|---|---|
| 12:40308546:A:G | K1374E | 7.23e-6 | MCPS Variant browser |
| 12:40308547:A:G | K1374R | 6.20e-7 | gnomAD browser |
| 12:40308549:A:C | T1348P | 1.86e-6 | gnomAD browser |

*Table S1: LRRK2 GTP-binding variants identified in the MCPSV and gnomAD variant browsers.*
